# Non-Elastic Remodeling of the 3D Extracellular Matrix by Cell-Generated Forces

**DOI:** 10.1101/193458

**Authors:** Andrea Malandrino, Michael Mak, Xavier Trepat, Roger D. Kamm

## Abstract

The mechanical properties of the extracellular matrix (ECM) – a complex, 3D, fibrillar scaffold of cells in physiological environments – modulate cell behavior and can drive tissue morphogenesis, regeneration, and disease progression. For simplicity, it is often convenient to assume these properties to be time-invariant. In living systems, however, cells dynamically remodel the ECM and create time-dependent local environments. Here, we demonstrate that cell generated contractile forces are capable of producing substantial irreversible changes to the density and architecture of physiologically relevant ECMs – collagen I and fibrin – in a matter of minutes. We measure the 3D mechanical deformation profiles of the ECM surrounding cancer and endothelial cells during stages when force generation is active or inactive. We further correlate these measurements to both discrete fiber simulations that incorporate fiber crosslink unbinding kinetics and continuum-scale modeling. Our findings reveal that plasticity, as a mechanical law in these networks, is fundamentally related to the force-driven unbinding of fiber crosslinks. These results illustrate the dynamic nature of the mechanical environment of physiologically mimicking cell-in-gel systems.

## Introduction

The Extracellular Matrix (ECM) is a scaffolding medium that helps transmit mechanical signals among cells in cancer^1,2^, fibrosis^3,4^, vascular networks^5,6^, and more generally, morphogenesis^7,8^. The mechanical and biochemical properties of the ECM impact cell behavior. The stiffness of the local environment and the tensional response from cells can induce invasive phenotypes in tumors^9^–^12^, guide differentiation in stem cells^13^, and alter vascular function ^14^. The fibrillar nature and local architecture of the ECM can lead to directed cell migration ^15^, and increased density and alignment in the tumor stroma are correlated with more aggressive disease and worse prognosis in preclinical and clinical samples^16,17^. It is not clear, however, how ECMs in living systems evolve spatially and temporally to promote physiological and pathological states.

Many studies exist on the quantification of mechanical signals traveling across ECMs, mostly assuming ideal ECM material properties. Studies usually derive the magnitudes of forces exerted by cells through imaging of fluorescent markers tethered to the ECM^18,19^. Because it is difficult to back-calculate forces in heterogeneous, dynamic environments, these approaches rely on 3D biopolymers or 2D substrates with time-invariant mechanical responses. The spatiotemporal evolution of the ECM is however relevant in many mechanobiological processes^3,20,21^. For instance, in angiogenesis and vasculogenesis, together with chemical signaling driving formation or inhibition patterns^22^, endothelial cells mechanically sense each other^23^, and cooperate to form tubular shapes by remodeling the fibrous 3D ECM^5,6^. Furthermore, mechanical signals are amplified, resulting in long-range force transmission, when ECMs are fibrillar, via local alignment and force-driven anisotropy^4,24,25^. More complex descriptions of fibrous materials are taken into account in recent 3D studies of forces in biological processes^26^.

ECM remodeling remains very challenging to decipher, despite its biological ubiquity. Remodeling entails dynamic molecular processes such as cell-fiber interactions, proteolytic degradation, and crosslinking sites binding and unbinding that ultimately lead to global changes in the ECM network. Additionally, cell-scaled forces are sufficient to drive ECM remodeling, and remodeled ECMs can in turn modulate mechanosensing in cells, resulting in dynamic feedback. The dynamic mechanical states of cells and the ECM, especially in physiologically relevant conditions, are not well understood.

Here we investigate cell-induced remodeling of physiologically relevant 3D fibrillar ECMs, specifically fibrin and collagen. We focus on cellular force-induced non-reversible reassembly of the ECM, which interestingly occurs on the time scale of minutes and leads to drastically different local architectures. We perform computational simulations with a network model with discrete ECM fibers to examine quantitatively the impact of cell forces and reaction kinetics at the ECM component level on tension and concentration profiles across the ECM network. We ultimately propose that constitutive damage and plastic softening at the continuum level are capable of recapitulating both experimental and fiber-level simulation findings.

## Results

### Experiments

We explore remodeling and the interaction with crosslinking level in two combinations of cell-ECM types cultured in 3D conditions, to explore the remodeling in relevant organ and disease models. As a first combination, we use endothelial cells in fibrin gels given the well-known ability of these cells to form physiologically mimicking microvascular network structures^27^. As a second combination, we use breast cancer cells in collagen I gels, a highly abundant component in the tumor stromal microenvironment. We start the experiment under inhibition of cell-generated forces with Cytochalasin D to obtain a force-free configuration (Fig. 1a). By washing Cytochalasin D, we let cells recover the ability to generate forces over a period of several hours. After recovery, we perform a cell lysis (decellularization) with a detergent to obtain the final force relaxation. We quantify deformations with a 3D Digital Volume Correlation (DVC) algorithm^28,29^ coupled with fluorescently labeled fibrin and collagen gels. In all cell-ECM combinations, remodeling involves non-elastic, *i.e.* non-recoverable, deformations of fibers. This plastic remodeling of recruited fibers is force-driven; it does not occur when Cytochalasin D is present and cell-generated forces are inhibited (Fig. 1a). Increasing crosslinking results in the decrease of bulk displacements (Fig. 1b), consistent with reports of more than two-fold increase of bulk stiffness, and ten-fold increase in the single filament stiffness going from unligated (low crosslinking) to ligated (high crosslinking) fibrin gels^30^. Crosslinking also affects the plastic remodeling, with poor crosslinking resulting in the enhancement of fiber densification at the cell-matrix boundary (Fig 1a,c). We further define the recoverability index as the ratio between the recoverable elastic deformation after decellularization and the maximum overall deformation—elastic and plastic—occurring over the course of experiment. For all cases, after release of the inhibitor of contractility, Cytochalasin D, we observe a significant remodeling in less than 1 hour (Fig. 1b, c, d). To dissect apart the effect of recovery from Cytochalasin D, we also perform experiments and displacement calculations immediately after seeding (without Cytochalasin D) and demonstrate that the remodeling dynamics occur over the course of minutes and the remodeling rate diminishes within an hour (Fig. 1d, Supplementary Fig. 1, and Supplementary Videos 1 and 2). All cell-ECM combinations here display plateauing cumulative remodeling in the course of hours with substantial irreversible components (Fig 1b and Supplementary Fig. 2). For fibrin, lower crosslinking leads to a lower recoverability index (Fig. 1e).

**Figure 1:**
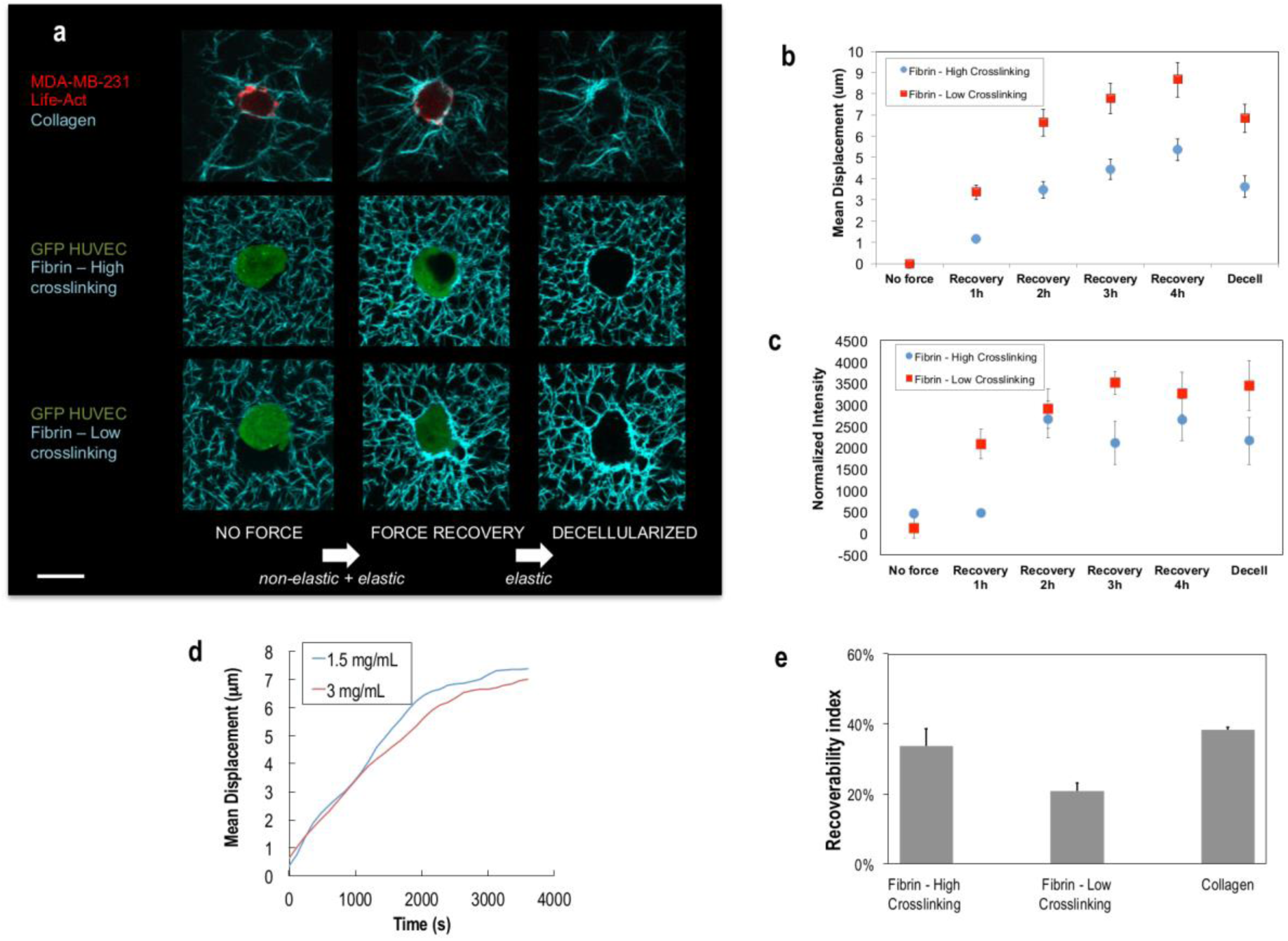
Non-elastic remodeling occurs in the course of minutes and depends on crosslinking in 3D biopolymer networks. **a**, Z-projected confocal images of human cells including HUVECs and MDA-MB-231 Life-Act line suspended in 3D biopolymer networks of fibrin (3 mg/mL) and collagen (1.5 mg/mL), respectively. For fibrin gels, crosslinking was lowered using a transglutaminase inhibitor (DDITS, 0.2 mM). White arrows indicate the assumption of the type of deformation, overall –elastic *and* non-elastic– or elastic only, expected between two force configurations. Scale bar, 20 µm. **b**, **c**, Quantification of the effect of crosslinking in terms of mean total displacement from the FIDVC algorithm inside a 60 µm ROI (region of interest), each containing one cell (N=8 cells) (b), and in terms of concentration changes (e.g. fiber recruitment) radially within 10µm from the cell membrane (c). Recovery is defined as the time after Cytochalasin D is removed. Concentration changes are expressed as a normalized intensity value obtained from the Z-projected confocal images (N=8 cells). **d**, Representative ECM remodeling dynamics by MDA-MB-231 cells for two different collagen gel densities. To assess the dynamics without the force recovery effect, Cytochalasin D is not used in these experiments. The mean cumulative matrix displacement in a ROI (∼30x30 µm^2^) containing the cell and the newly recruited ECM fibers is obtained starting at the reference configuration shortly after seeding. These relatively fast dynamics are calculated from time-lapse images of projected z-stacks. **e**, Summary of the Recoverability Index for all the cell-matrix pairs studied.

We further consider how cell-remodeled ECMs transmit forces in space. Both endothelial and cancer cells apply centripetal tractions, as demonstrated by the directions of the local matrix displacements (Fig. 2a,c). Plastic recruitment leads to a substantially higher magnitude of cumulative matrix displacements than the elastic displacement magnitude alone (Fig. 2c). By normalizing displacement propagation through the ECM, we discover that long-range force transmission depends strongly on the specific cell-ECM pair, with the displacements decaying steeper in collagen than fibrin. In all cases, spatial displacement decays slower than in the ideal case of isotropic linear elastic material, confirming long-range mechanical signaling capability (Fig. 2b). Fully and partially crosslinked fibrin matrices behave similarly in their ability to propagate displacements (Fig 2b, solid lines). We also observe similarities when quantifying how the displacements generated by cell contraction during remodeling decays (overall displacement), as compared to how the elastic component of the displacement decays (based on measurements right before and after decellularization and relaxation).

**Figure 2:**
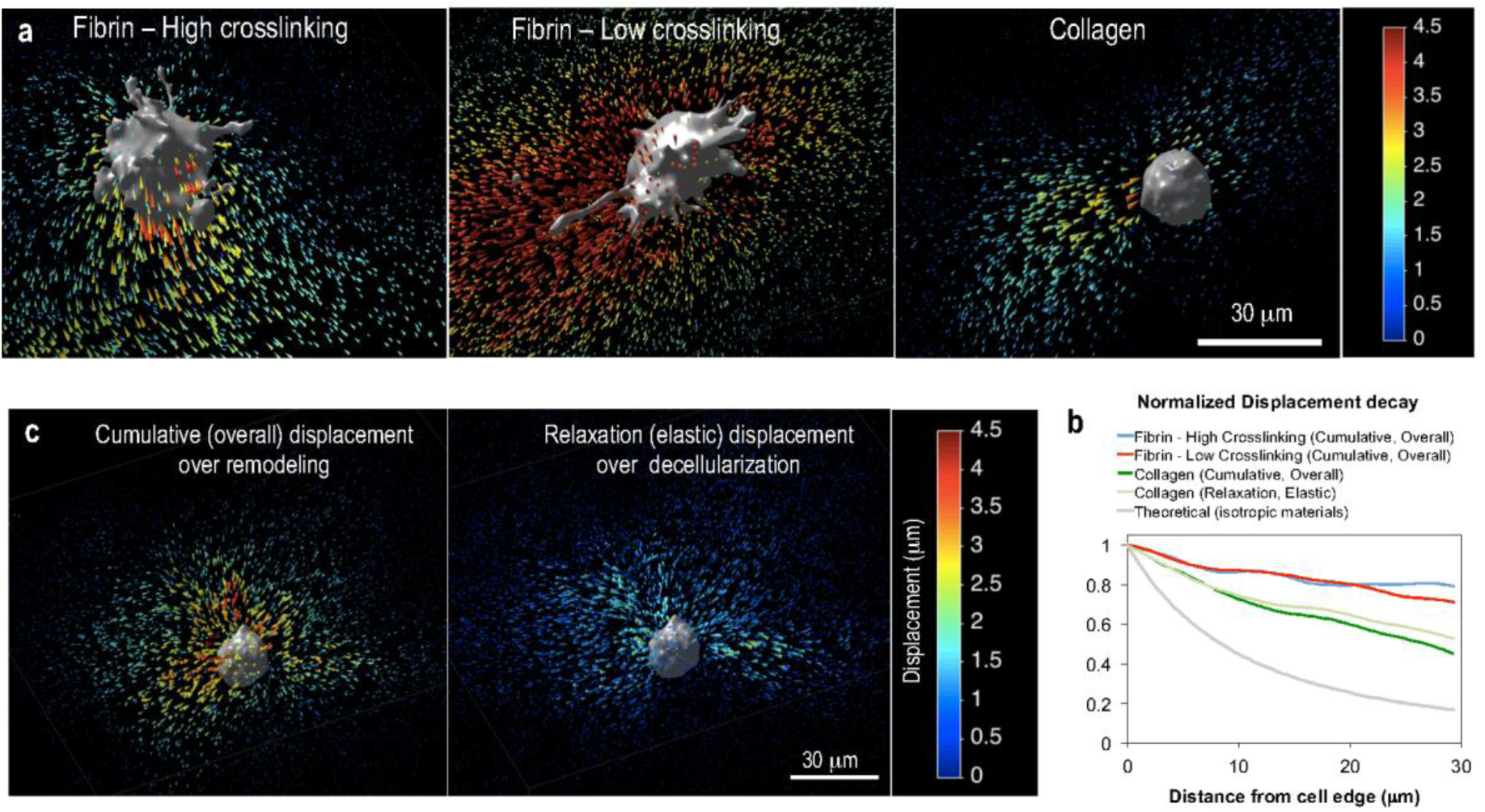
Mechanical signaling through long-range displacement propagation is cell-matrix specific and can be modified by remodeling. **a**, 3D displacement vectors from the FIDVC algorithm, color-coded according to the magnitude of the cumulative displacement, for three representative cells in the three different matrices tested. Cumulative displacement includes both elastic and plastic components over the duration of the ECM remodeling process. **b,** Normalized displacement decays versus distance from the cell membrane for the matrix cases analyzed (N=5 cells per condition). For a given cell, the decay was calculated by averaging from four in-plane, radially outward lines. A theoretical derivation for isotropic materials, decaying as (distance)^-2^, is shown for comparison. **c,** 3D displacement field for a representative case of a breast cancer cell in collagen over two stages: (left) force recovery with matrix remodeling, with cumulative displacements being both plastic and elastic, and (right) force relaxation over decellularization, with only the elastic component of the displacements expected. Relaxation displacements are shown in the reverse direction to display the centripetal elastic displacements that the cell is inducing right before being lysed.

Questions remain regarding the mechanics and dynamics of cell-ECM interactions in complex 3D matrices. In previous studies, confocal reflectance imaging has been used as a label-free technique to visualize and characterize heterogeneities in collagen matrices under cellular forces^26,31^. Anisotropic or isotropic elastic models used in these investigations, however, do not provide insights about non-elastic remodeling by cell-generated forces. Recently, in a study using 3D DVC algorithms and fluorescently-labeled fibrin matrices, Notbohm *et al.* characterized the remodeling process and inferred that fibroblasts plastically push and pull the fibrin to form tubular, protrusion-like permanent structures^32^. Other assessments using fluorescently labeled fibers in experiments have focused on continuum properties and characterizing the viscoplasticity of the ECM in reconstituted and living microtissues^33,34^. While useful and informative, these studies do not provide experimental assessment of the effects of varying crosslinking concentrations. Also, while the molecular-level mechanistic insight was recognized, a multiscale computational approach linking discrete fiber networks with local force-sensitive crosslink kinetics to a coarse-grained continuum representation of the ECM has not be fully investigated. We also confirm the mechanical signaling enhancement provided by ECMs with fibers as compared to isotropic matrices. This has been previously attributed to non-linear phenomena such as strain stiffening^23^, or local alignment by cell-generated forces – a load-driven anisotropic effect in elastic matrices^24,25^. Here, we additionally show that such signaling can change dynamically because the matrix is plastically remodeled. These results point to the need for more investigations on the possible role of force-induced ECM remodeling in cell mechano-sensing dynamics in physiological 3D environments.

### Simulations

We perform computational simulations to extract quantitative details of how local molecular-level features can influence global reorganization dynamics of the ECM under cell-generated forces. A 3D fiber network, mimicking the ECM, is generated by polymerizing monomeric units into elastic fibers that can stretch and bend, according to the following potentials:

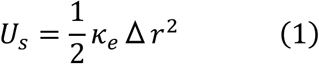

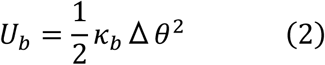

where *U_s_* is the potential energy from stretching, *U_b_* is the potential energy from bending, *κ_e_* is the extensional stiffness, *κ_b_* is the bending stiffness, Δ*r* is the deviation from the equilibrium length, and Δ*θ* is the deviation from the equilibrium angle. Each monomeric unit adds a cylindrical segment to the fiber during polymerization, and fibers nucleate in random directions during initial network formation. Proximal fibers are connected with crosslinks that can unbind in a force-sensitive manner in accordance to Bell’s model ^35^

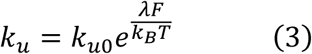

where *k_u_* is the crosslink unbinding rate, *k*_*u*0_ is the zero-force unbinding rate, *λ* is the mechanical compliance of the crosslink, *F* is the magnitude of the extensional force acting on the crosslink (for positive stretch), *k_*B*_* is the Boltzmann constant, and *T* is temperature. Model parameters are listed in Supplementary Table 1. Parameter values for the simulated fibers and network are chosen based on plausible values for fibrin ^30,36^ and experimental network features (Supplementary Fig. 3). We note here that ECM fibers are typically complex, hierarchical structures. For example, during gelation, fibrin molecules polymerize into fibrils that bundle into thicker fibers. Thicker fibers can be linked to other fibers, forming a connected network ^37,38^. It is possible that mechanical forces can disrupt both inter-fiber and intra-fiber bonds. For simplicity and computational feasibility, we only consider the thick fiber structures and one type of crosslink (which connects different fibers). This mimics reasonably closely our *in vitro* experiments where fiber-fiber connections are prominent.

**Figure 3:**
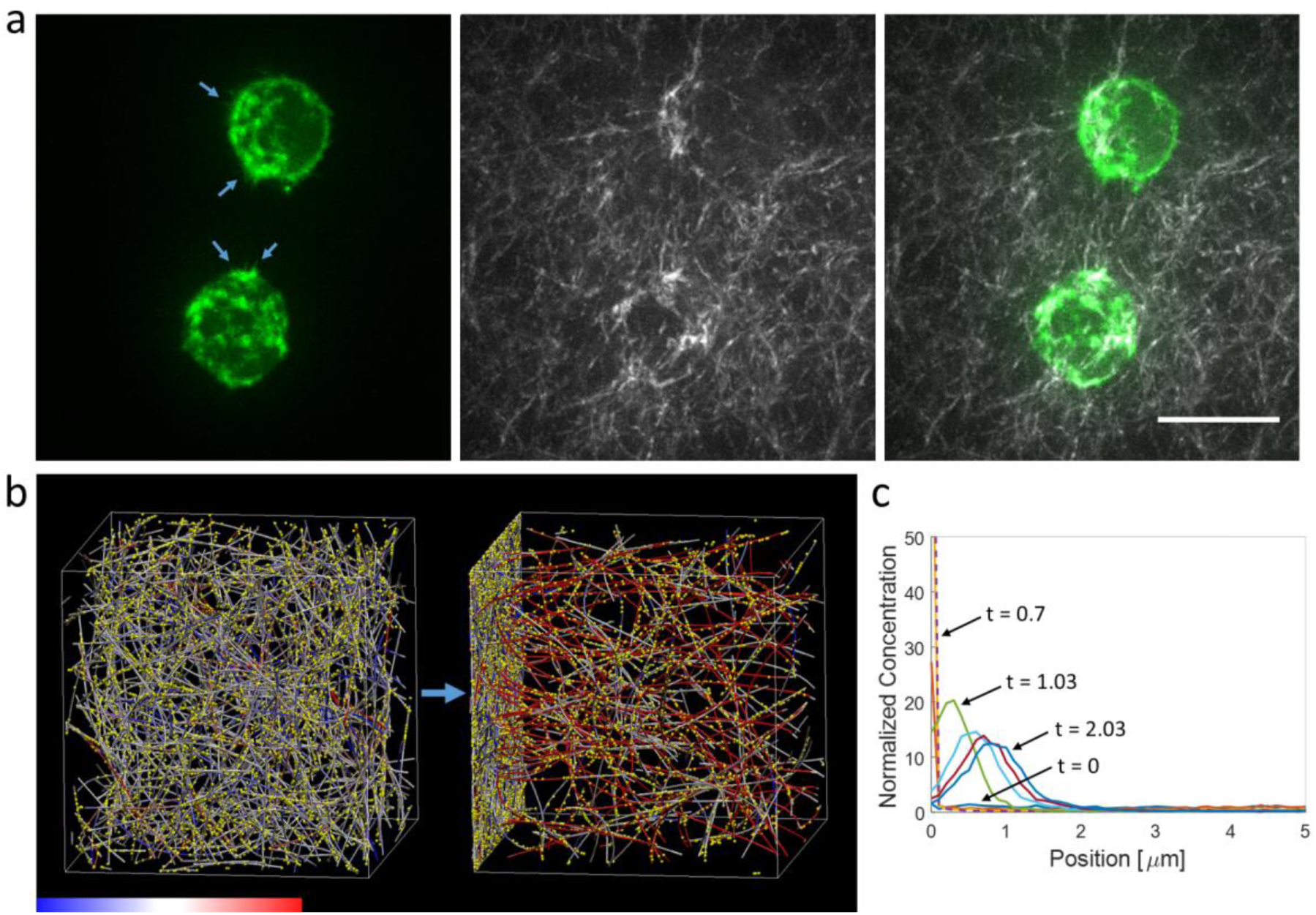
Actin dynamics and ECM recruitment. **a**, MDA-MB-231 cells expressing fluorescent F-actin (green, left) inside a 3D collagen matrix (white, middle) with a concentration of 1.5mg/mL display many dynamic actin protrusions (blue arrows). Overlay image of actin and collagen is on the right. Images are maximum intensity z-stack projections. The scale bar is 20µm. See also Supplementary Video 3. **b**, Computational simulation of an ECM fiber network shows network morphology before (left) and after (right) the application of loading forces near the left boundary. Colors on fibers indicate tension level according to the color bar (-300 to 300pN). Yellow spots are crosslinks (places where fiber-fiber crosslinking can occur). See also Supplementary Videos 4 and 5. **c,** Overlay of the time evolution of fiber concentration profiles, normalized by the initial concentration, in the force-loading direction as loading forces are exerted from the left boundary. Loading forces mimic dynamic filopodia pulling from the loading boundary, such that fibers within 2µm of that boundary exhibit a force pulling them toward the boundary. As new fibers or fiber segments move within that distance, new loading forces are exerted on them. Different colored curves represent different normalized times of: 0 (lower blue, uniform), 0.03 (red), 0.37 (yellow), 0.7 (purple, dashed), 1.03 (green), 1.37 (light blue), 1.7 (magenta), 2.03 (blue), where time is normalized to the total time of force application (starting at right after 0 and ending at 1). Some relaxation occurs after applied forces end, but fiber recruitment here is not reversed. The simulation setup is 100pN loading per fiber segment in the loading region, 1x crosslink zero-force unbinding rate, 0.3x crosslink mechanical compliance, and 1x crosslink density (see Supplementary Table 1 for default values).

Once the crosslinked network is generated and relaxed to a stable state, a loading force is applied on one side of the network for a fixed duration of time and then reduced to zero to allow the network to relax to a new, potentially remodeled state. The force loading is applied such that any fiber segment that reaches within a certain distance (2µm) of the –z boundary experiences a local point force pulling it toward that boundary. We explore a range of force magnitudes from 1pN to 1nN to capture the impact of physiologically plausible cell generated forces. The loading condition mimics a pulling process where new filopodia are continuously generated that adhere to and pull new fiber segments near the cell. This type of loading is needed in order for fibers to be continuously recruited toward the cell, and many dynamic actin protrusions are indeed observed on the periphery of cells embedded inside a 3D ECM as shown in Fig. 3a and Supplementary Video 3. The fiber ends at the +z boundary are fixed to mimic the resistance from fibers far away. The x and y boundaries are periodic, and the domain size is 20x20x20µm^3^. The network is athermal and the components (fiber segments, crosslinks) follow the equation of motion:

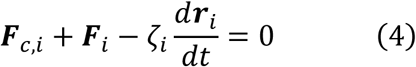

where *i* is the index of the component under consideration, ***F**_c,i_* is the cell generated loading force near the –z boundary, ***F**_i_* is the mechanical force from the fiber network, which includes extension, bending, and repulsion of the fibers and crosslinks, *ζ_i_* is the drag coefficient, and ***r**_i_* is the position. Sample simulation network and dynamics are shown in Figs. 3b,c and Supplementary Videos 4 and 5. Additional details of the discrete fiber network model, which has been applied previously to simulate other filamentous networks, can be found here ^39,40^.

**Figure 4:**
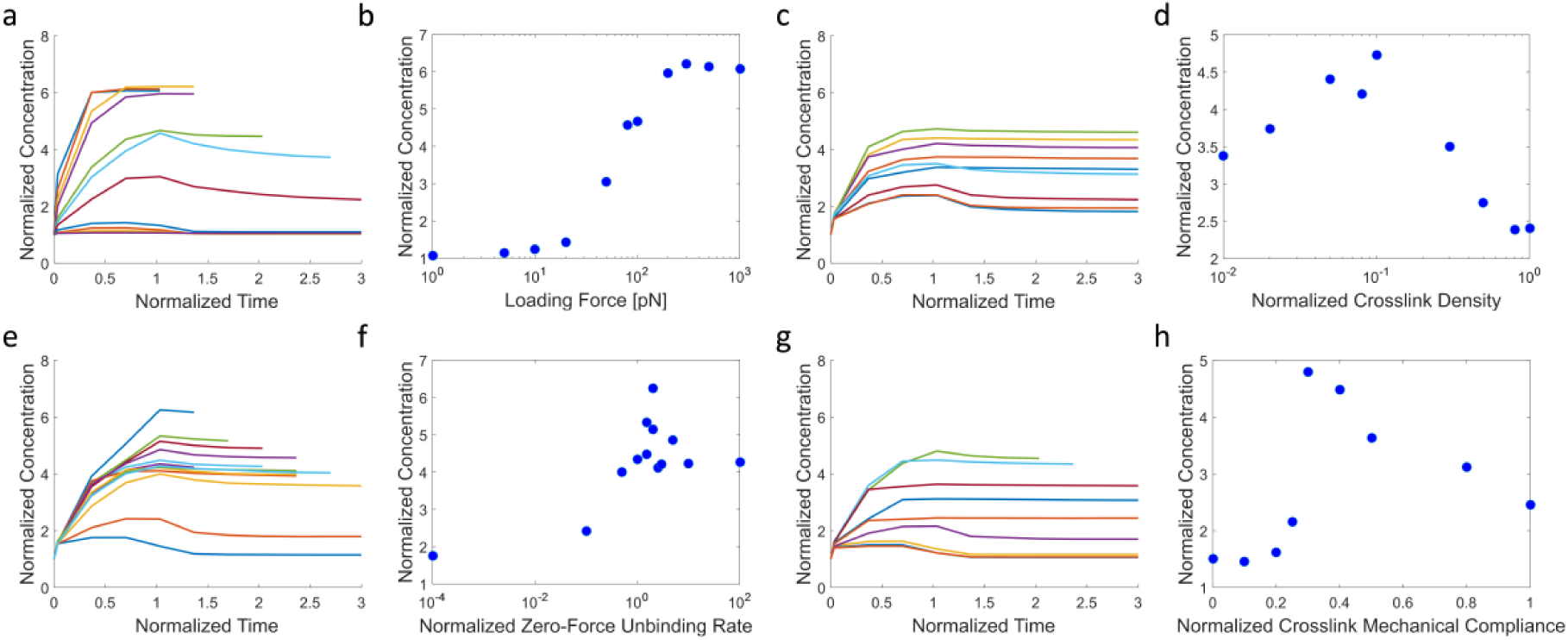
Filament dynamics analyses reveals biphasic dependence of fiber recruitment on both crosslink density and compliance and plateauing dependence on both loading force and unbinding rate. **a**, Different colors represent loading forces (per fiber segment in the active loading region) of 1000, 500, 300, 200, 100, 80, 50, 20, 10, 5, and 1 pN, for blue, red, yellow, purple, green, light blue, magenta, lower blue, and lower red curves, respectively. Time is normalized such that the loading forces are stopped at 1, which is also approximately the time when fiber recruitment has reached near the peak. The normalized concentration is that within 2.5µm from the loading boundary (accumulation region) divided by the initial uniform concentration before applied forces. **b**, Maximum normalized concentration in the accumulation region as a function of the loading force. The crosslink zero-force unbinding rate is 1x, the crosslink mechanical compliance is 0.3x, and the crosslink density is 1x. **c**, Different colors represent different relative crosslink densities of 0.01x, 0.02x, 0.05x, 0.08x 0.1x, 0.3x, 0.5x, 0.8x and 1x for blue, red, yellow, purple, green, light blue, magenta, lower blue, and lower red, respectively. **d**, Maximum concentration in the accumulation region as a function of the normalized crosslink density. The loading force is 100pN, the crosslink zero-force unbinding rate is 0.1x, and the crosslink mechanical compliance is 0.3x. **e**, Different colors represent networks with different crosslink zero-force unbinding rates of 0.0001x, 0.1x, 0.5x, 1x, 1.5x, 1.5x, 2x, 2x, 2.5x, 3x, 5x, 10x, and 100x for blue, red, yellow, purple, green, light blue, magenta, upper blue, and upper red, upper yellow, upper purple, lower green, lower light blue, respectively. **f**, Maximum concentration in the accumulation region as a function of the crosslink zero-force unbinding rate. The loading force is 100pN, the crosslink density is 1x, and the crosslink mechanical compliance is 0.3x. **g**, Different colors represent networks with crosslink mechanical compliance of 0, 0.1x, 0.2x, 0.25x, 0.3x, 0.4x, 0.5x, 0.8x, and 1.0x for blue (lower), red (lower), yellow, purple, green, light blue, magenta curves, blue (higher), and red (higher) respectively. **h**, Maximum concentration in the accumulation region as a function of mechanical compliance of crosslinks. The loading force is 100pN, the crosslink zero-force unbinding rate is 1x, and the crosslink density is 1x. See Supplementary Table 1 for 1x values of the parameters.

We examine the network remodeling dynamics under different conditions, modulating experimentally tunable or physiologically relevant parameters. Specifically, we consider different loading forces, crosslink densities, zero-force unbinding rates of crosslinks, and crosslink bond mechanical compliances. These parameters aim to capture the impact of cell traction, the degree of ECM crosslinking, and the kinetic nature of crosslinks. As shown in Fig. 3, ECM fibers are recruited to the loading boundary once applied forces are activated. Temporal profiles of the ECM concentration near the loading boundary during and after forces are applied under the different conditions are shown in Figs. 4 a, c, e, and g. In some conditions, after the force is deactivated (at a normalized time of 1), the network primarily recovers elastically and the normalized ECM concentration near the loading boundary returns toward 1, equivalent to the uniform network state prior to loading. The conditions leading to this behavior are low applied forces, high crosslink density, low zero-force crosslink unbinding rate, and low crosslink mechanical compliance. This indicates that weak cells and strongly crosslinked networks exhibit low plastic remodeling in the ECM. Conversely, plastic remodeling, in which the recruited ECM fibers do not relax back to their original positions after force loading is stopped, is exhibited under high loading forces and networks with weaker crosslinking. This is demonstrated in Supplementary Videos 4 and 5 and Fig. 3c, which show fiber recruitment due to loading forces and the network permanently remodeled with recruited fibers remaining near the loading boundary after the cessation of applied forces. There also tends to be a biphasic relationship between fiber recruitment and the parameters explored, as shown in Figs. 4 b, d, f, and h. Fiber recruitment vs. loading force (log scale) displays a sigmoidal trend, under which minimal ECM fiber recruitment occurs under low loading forces below a threshold, increasing ECM fiber recruitment occurs with increasing loading forces at an intermediate range, and reaching a plateau for forces above a second threshold. Under the same loading forces, increases in crosslink density first lead to more ECM recruitment, but as the crosslink concentration increases beyond a certain threshold, network remodeling is reduced. This results from the network gaining connectivity with higher crosslink concentration, enabling connected fibers farther away to be recruited, but beyond a certain concentration the loading force is distributed between more crosslinks leading to reduced crosslink unbinding rates. Similarly, for altered zero-force unbinding or mechanical compliance of the crosslinks, crosslinks that do not unbind easily lead to networks that are not remodeled significantly and have low fiber recruitment. Crosslinks that unbind at high rates or with high force sensitivity lead to networks that undergo loss of connectivity such that fibers farther away from the loading boundary are not effectively recruited.

We further consider the overall stress profiles in the network under our loading condition. Stresses are calculated by summing the normal component of forces exhibited by fibers crossing a plane parallel to the plane of the loading boundary divided by the area of the plane. As shown in Fig. 5, the stress profiles during the loading period (the normalized time range from 0 to 1) tend to be dynamic, especially when fibers are being recruited plastically. Under varying loading force magnitudes (Fig. 5a), for high forces, the overall stress on the network peaks and then decays as fiber crosslinks unbind leading to relaxation. At relatively low applied forces, the stress profile does not decay and instead reaches a plateau value, as crosslink unbinding and network relaxation does not occur during the loading period. When the crosslink density is varied under the same loading forces (Fig. 5b), the network stress is low at low crosslink density as unbinding prevents the build-up of stress, and the network stress is high and sustained a high crosslink density as stresses can build up and maintain with low crosslink unbinding. At the intermediate level, stresses can build up, due to sufficient crosslinks present, followed by relaxation due to crosslink unbinding. Similarly, when the kinetics of the crosslinks is tuned (Figs. 5 c and d), more stable, permanent crosslinks lead to large stress build-up and sustained stress levels, while crosslinks that unbind more quickly lead to reduced and less sustainable stresses. Moreover, crosslinks that unbind more quickly, either with a higher zero-force unbinding rate or higher mechanical compliance, lead to stress profiles that decay more quickly over time. This indicates a scaling relation between the overall dynamics of ECM stresses (and fiber recruitment) with crosslink density and kinetics. Furthermore, the simulations discussed so far do not consider the possibility of the rebinding of unbound crosslinks. We find that enabling rebinding appears to partially diminish ECM recruitment and network stress dissipation (Supplementary Figure 4).

**Figure 5:**
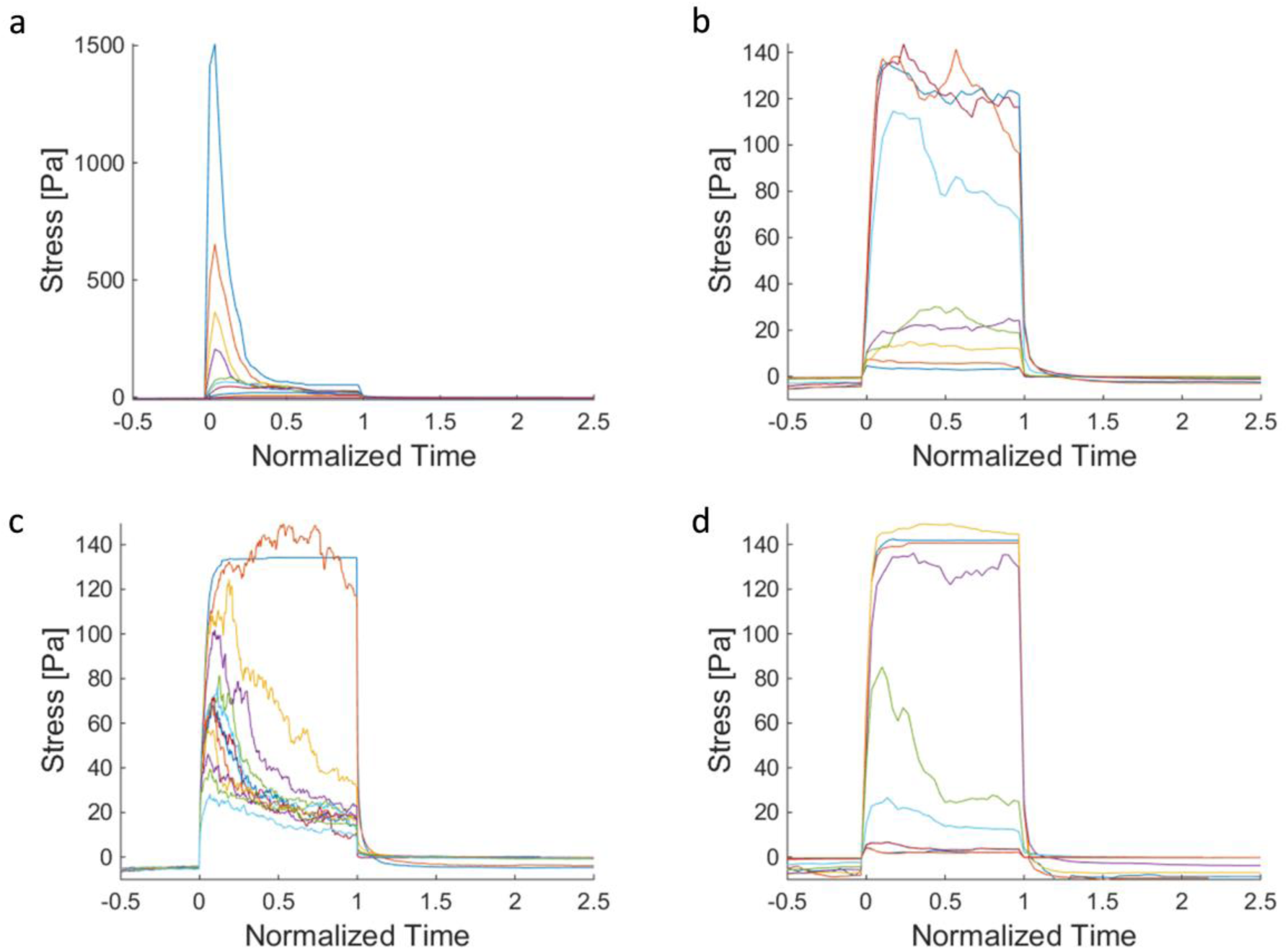
Stress generation in the ECM networks highly depends on remodeling of ECM filaments. Stress vs. time plots for ECM networks during loading. Force loading starts at 0 and stops at the normalized time of 1. The stress level in the network initially rises and peaks as loading is initiated. Then, as the crosslinks unbind, the stress relaxes. **a**, Stress vs. loading force; top to bottom (high to low loading forces); **b**, vs. crosslink density; bottom to top (low to high crosslink density); **c**, vs. crosslink zero-force unbinding rate; top to bottom (low to high zero-force unbinding rate). **d**, vs. crosslink mechanical compliance; top to bottom (low to high compliance). Coloring schemes for (a-d) are identical to Figs. 3 (a, c, e, g), respectively.

**Figure 6:**
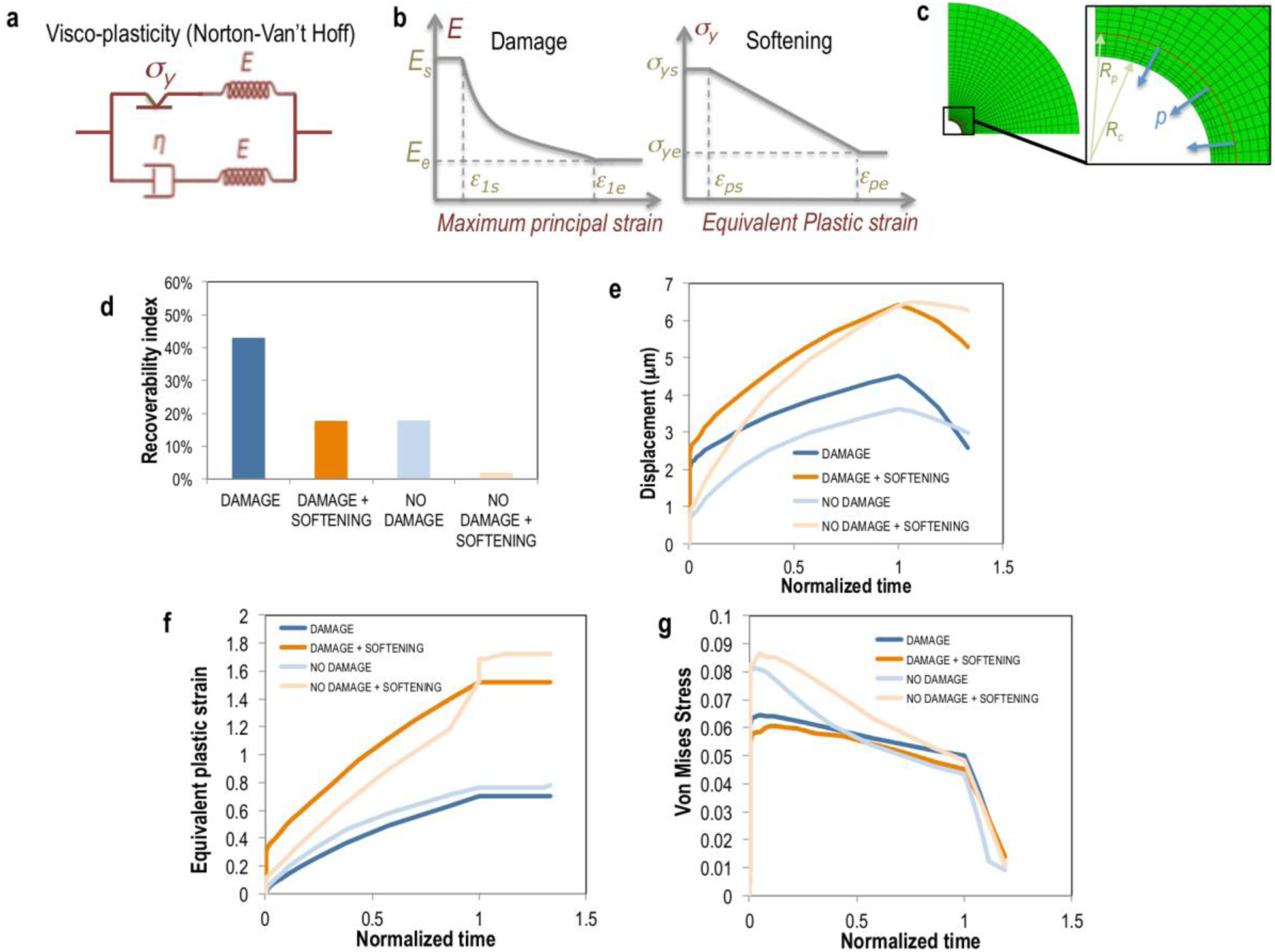
A strain-dependent plastic softening with elastic damage constitutive law recapitulates the effect of crosslinking in fibrous matrices. **a**, Schematic representation of the constitutive viscoplasticity of the Norton Van’t Hoff type used to model the ECM. (Parameters provided in Supplementary Table 2.) **b**, The model is modified and alternatively tested to include phenomenologically the features of damage, simulating breakage of crosslinks with tensile strain, and plastic softening, simulating the drop in yield stress with plastic strain. **c**, The constitutive model is implemented in the commercial finite element solver ABAQUS using existing standard viscoplastic material models and custom subroutine implementation for the damage model. An axisymmetric mesh is shown to model the ECM around a contracting cell. The zoom inset shows the surface where the load is applied to simulate the action of filopodia farther away from the edge. **d**, Recoverability index at the cell-ECM interface, defined previously as elastic deformation divided by the total deformation for the cases with or without damage and softening. **e, f, g**, Displacement (**e**), plastic equivalent strain (**f**), and Von Mises equivalent stress (**g**) for the cases with or without damage and softening, all calculated at the edge cell-ECM. Loading starts at 0 and stops at the normalized time of 1.

We explore the possibility to coarse-grain the unbinding dynamics effect seen at the fiber level up to the material continuum level. Driven by creep and inelasticity observation in both our experiments and filament computer model results, we start from a constitutive model based on Norton-Hoff viscoplasticity (Fig. 6a and Supplementary Note): the viscous element simulates the creep response, and the plastic element simulates permanent, inelastic deformations. To recapitulate crosslink unbinding, we build up from this baseline viscoplasticity, by adding both a ‘damage’ feature, *i.e.* a decay in elastic stiffness starting above a critical strain, and a linear plastic softening, *i.e.* a decay of yield stress above a critical plastic strain (Fig. 6b). We implement and test the viscoplastic model with damage and softening features in a finite element simulation with spherical symmetry of a cell contracting centripetally and displacing similarly as in the experiments, and then relaxing the force; this force being applied slightly outward from the edge, simulating the action of filopodia recruiting relatively close fibers (Fig, 6c). This force is meant to produce mostly tensile stresses but also some degree of local compression as the material is recruited and packed at the cell-ECM interface. The presence of damage recapitulates (i) the increase in bulk displacement at the edge with long-term loading (Fig. 6e) observed when lowering the density of crosslinks (Fig. 1b); and (ii) the decrease in equivalent stress at the edge (Fig. 6g) as observed in the filament model when lowering the density of crosslinks (Fig. 5b). However, the damage feature is not enough to predict correctly the trend in recoverability index observed in the experiments (Fig 1e). Instead, plastic softening is needed to reproduce the loss in elastic recovery as density of crosslinks decreases (Fig. 6d) and the coupled increase in plastic strain (Fig. 6f).

These results at the material continuum level suggest that damage, simulating the unbinding and elastic failure of crosslinks, and softening, simulating the drop in average material yield, as fewer crosslinks are present, might act synergistically and be both necessary to model plastic remodeling of the ECM. We thus suggest that more experimental assessment is needed to unravel the role of viscoplastic damage and softening. Continuum relationships of this kind are important because they can retrieve more precisely values of tractions calculated from the experiments in fibrous and remodeled biopolymers. ECM viscoplasticity has been recently targeted for accurate descriptions of tissue dynamics^33^. However, to our knowledge, this is the first study that integrates experimentally tuned molecular features such as the crosslink density, with fiber-level force-driven unbinding dynamics, at a continuum-level to provide a mechanistic view of ECM plastic remodeling.

Overall, our results suggest that during ECM recruitment, cells do not exhibit a stable tensional state, but rather a highly dynamic one due to relaxation from crosslink unbinding. Thus, for highly motile cells that recruit matrix, including endothelial cells and metastatic cancer cells, as they migrate to and recruit fibers from new locations inside a 3D ECM, their tensional profile is dynamic rather than static. This is a starkly different picture compared to cells on 2D artificial elastic substrates, as those cells tend to exhibit relatively stable stress profiles since the substrates do not undergo remodeling ^41^. Many studies have shown that substrate stiffness affects cell tension and this, in turn, affects cell behavior. However, the dynamic state of cell tension, which appears to be characteristic of cells inside a more physiologically relevant environment of a 3D ECM, has not been fully investigated. Our study shows that certain key properties – cell loading forces, ECM crosslink density, and the kinetic nature of the crosslinks – are physically implicated in regulating this dynamic tensional state in cells, which can help guide future experiments in systematically tuning these parameters to assess cell behavior.

## Conclusion

The physiological microenvironment is often composed of a complex, fibrillar ECM that exhibits non-linear, non-elastic properties. We have demonstrated that cell-generated forces via the actomyosin machinery are capable of mechanically reassembling the local ECM, leading to substantially increased local ECM density in the course of minutes, which is not fully reversible when the cells are relaxed. Differences in ECM ligand density can alter cell signaling and overall phenotypes^42^–^44^. The results demonstrated here highlight the dynamics of cell-ECM interactions in a more physiological context. The local environment sensed by cells, both physically and biochemically, is highly distinct from acellular matrices and gels in their initial states, with nominal concentration values based on stock solutions. ECMs with active cells are rapidly remodeled by cells to generate heterogeneous local environments with significantly different ligand densities and architectures. This behavior is often not considered, as only nominal ECM concentrations are usually reported, and is not captured by widely used non-physiological, elastic substrates that cannot be plastically remodeled by cells. Modified microenvironments have already been shown to lead to diverse ramifications in cell behavior, from guiding stem cell differentiation to modulating tumor dissemination and tissue morphogenesis. Our results directly implicate cell mechanics – the actomyosin machinery – in driving active remodeling of the ECM and the creation of new microenvironments that can dynamically modulate cell behavior.

## Acknowledgements

We thank Christian Frank and Mohak Patel from Brown University for their initial support in running the FIDVC code, members of Pere Roca-Cusachs and Trepat Lab for discussions. A.M. received funding from the People Programme (Marie Curie Actions) of the European Union’s Seventh Framework Programme FP7/2007-2013/under REA (Grant 625500). Funding from the U.S. National Cancer Institute (U01 CA202177-01) to R.K. is gratefully acknowledged. X.T. acknowledges the Spanish Ministry of Economy and Competitiveness (BFU2011-23111), the Generalitat de Catalunya (Programa CERCA and 2014-SGR-927), and the European Research Council (CoG-616480).

## Author Contributions

A.M., M.M., and R.D.K. designed the experiments, A.M. fabricated the microfluidic devices, A.M. and M.M. performed the experiments, M.M. performed the discrete model simulations, A.M. performed the continuum model simulations, and A.M. and M.M. wrote the manuscript. A.M., M.M., R.D.K., and X.T. analyzed the data, discussed and interpreted the results, provided technology, and commented on the manuscript.

## Methods

### Cell culture

For fibrin experiments, we culture Human Umbilical Vein Endothelial Cells (HUVEC) (Lonza) were cultured on collagen I-coated flasks in EGM-2 (Lonza) growth medium and used in experiments between passages 6–8. For collagen experiments, we also culture MDA-MB-231 cells expressing fluorescent actin filaments (LifeAct, gift from the Lauffenburger Lab) were cultured at 37°C, 5% CO2 with DMEM supplemented with 10% fetal bovine serum and 1% penicillin-streptomycin.

### Gel preparation and encapsulation

Both collagen (Corning) and fibrinogen proteins (Sigma) are fluorescently labeled in stock solutions. A fluorescent reactive dye binding to free amine groups (NHS Succinimidyl Ester (Thermo Fisher) is used to produce cell-compatible, purified gels that can be visualized in 3D confocal live imaging. Stocks solutions are purified from the unreacted dye by using dialysis cassettes (Thermo Fisher) with a 7 kDa molecular weight cut-off.

Fluorescently labeled fibrin is then obtained by mixing over ice (i) fibrinogen dissolved in PBS (Lonza) at twice the final concentration (6 mg/mL) and (ii) Thrombin (Sigma), dissolved at 2U/mL in EGM-2 growth medium with HUVECs. Briefly, HUVEC are spun down at 1200 rpm for 5 min and the cell pellet is resuspended in EGM-2 growth medium containing the thrombin and mixed with the fibrinogen solution at a 1:1 ratio. The mixture is quickly pipetted into the device using the gel filling ports. The device is placed in a humidified enclosure and allowed to polymerize at room temperature for 10 min before fresh growth medium is introduced before the experiment to hydrate the gel.

Collagen gels are prepared by mixing type I rat tail collagen (BD Biosciences) with a neutralizing solution (100mM HEPES buffer in 2X phosphate buffered saline at pH 7.3) at a 1:1 ratio and then diluting with 1X PBS and suspended cells in media to a final collagen concentration of 1.5 mg/mL^45^. The final solution is then allowed to gel in a humidified chamber at 37 °C and 5% CO^2^.

### 3D chambers

Microfluidic devices with gel and media chambers are used because of the convenient fluid flow access for on-stage media and reagent exchange necessary for the experiments. Device design and protocol are described previously^46^. Briefly, 130 µm thick microfluidic devices were fabricated using PDMS soft lithography. The chambers are 1.3 mm wide and are injected with the gel encapsulating cells. Similarly shaped chambers for media flank these gel chambers and allow the quick washing and re-introduction of small volumes of reagents in all stages of the experimental procedure.

### Experiment and displacement mapping

In most of cell encapsulation experiments, the gels are polymerized together with cells treated with Cytochalasin D (5 µM), which is an inhibitor of actin polymerization and leads to highly diminished cellular force generation^47^. First, images are captured of cells encapsulated in the 3D ECMs under the action of Cytochalasin D to have a force-free initial configuration. Second, the chambers are washed through the microfluidic channels with fresh media on-stage three times to remove the Cytochalasin D, and to observe the onset of ECM remodeling. During this process, fluorescently labeled fibers are imaged at small time increments and these sequential images are cross-correlated through the Fast Iterative Digital Volume Correlation (FIDVC) algorithm^29^ to determine the 3D displacement field while remodeling occurs. After plastic remodeling of the ECM begins to plateau, a non-ionic detergent (Triton X, 0.1%) that preserves the gel structure while permeabilizing the cell membrane is used to lyse cells. Thus, cellular forces are fully relaxed at the final fiber network configuration. To obtain an estimation of the remodeling dynamics without the delay from Cytochalasin D recovery, FIDVC-based displacement estimations are also performed on time-lapse videos of cells right after seeding. Spatially averaged values of total displacement in Fig. 1 are obtained from FIDVC results inside a cubic ROI of 60-µm lengths, with each ROI containing one cell.

